# Brain stimulation preferentially influences long-range projections

**DOI:** 10.1101/2025.02.19.639189

**Authors:** Pedro G. Vieira, Matthew R. Krause, Pooya Laamerad, Christopher C. Pack

**Author notes:** These authors contributed equally (Pedro G. Vieira, Matthew R. Krause). Perelman School of Medicine, University of Pennsylvania (Pooya Laamerad).

## Abstract

Advances in brain stimulation have made it possible to target smaller and smaller regions for electromagnetic stimulation, in the hopes of producing increasingly focal neural effects. However, the brain is extensively interconnected, and the neurons comprising those connections may themselves be particularly susceptible to neurostimulation. Here, we test this hypothesis by identifying long-range projections in single-unit recordings from non-human primates receiving transcranial alternating current stimulation (tACS). We find that putative long-range projections are more strongly affected by stimulation than other cells. Specifically, they are both more entrained on average and account for occurrences of extremely strong entrainment. Since stimulation appears to target the edges, rather than nodes, of neural networks, it may be necessary to rethink how neurostimulation strategies are designed.

## Introduction

The discovery of brain regions specialized for specific behavioral functions or mental states [1–4] has inspired the development of methods for electromagnetic brain stimulation. These methods often aim to selectively target one brain region, thought to be responsible for particular behaviors or disease symptoms, in order restore or investigate its normal function.

Technical advances have now made it possible to generate increasingly focal electromagnetic fields inside the brain [5–9]. This is especially true for transcranial electrical stimulation (tES), a non-invasive method which delivers current through electrodes attached to the scalp [10]. Early tES experiments large sponges to produce electric fields that covered a target—but also much of the hemisphere around it. The development of multiple electrode systems, placed in personalized montages, can produce electric fields that are seven times more focal [9] and further improvements are expected by using interferential stimulation [8,11].

These advances are predicated on the idea that more focal electromagnetic fields will produce more focal neural effects, which in turn will permit more precise control over behavior or symptoms. However, this assumption remains to be tested. The interconnected nature of the brain allows even very local effects to propagate to distant sites through long-range connections. Moreover, the neurons that form these projections tend to be larger and may thus be more susceptible to the weak electric fields produced by tES. Thus, the focality of the stimulation itself may be distinct from the focality of its neural effects.

Understanding how stimulation affects long-range projections is therefore important for the future development of neuromodulation. To the extent that brain stimulation has primarily local effects, efforts to develop more focal brain stimulation methods become critical [8] and stimulation should be applied directly to the intended target. However, if stimulation preferentially targets long-range projections, this suggests the need for very different strategies based on the inputs and outputs of the target.

We have therefore examined these issues using single-unit recordings from alert non-human primates, a model organism that provides a close match to the biophysical and anatomical properties of the human brain. We first show that long-range projections can be identified on the basis of their extracellular waveforms and validate this approach pharmacologically. Next, we show that these putative projections are particularly strongly affected by electrical stimulation.

## Results

Our dataset comprised 428 neurons from four non-human primates (3 male, 1 female) and five brain areas (Area 7A, MT, V4, basal ganglia, and hippocampus). These data came from previous publications [12–15]. In each experiment, monkeys performed a simple visual fixation task meant to control fluctuations in endogenous brain activity due to changes in arousal, eye movements, and sensory stimulation. During this task, we applied blocks of transcranial alternating current stimulation (tACS), a form of tES that uses sinusoidal currents. These were interleaved with periods of sham stimulation (see Methods) or baseline measurement, allowing us to assess the effects of stimulation. To mimic human neuromodulation, tACS was applied to the monkeys’ intact scalps, using electrodes, equipment, and parameters designed for human use. Throughout this task, we recorded single-unit activity from sites within the focus of stimulation, as predicted by finite-element modelling based on each animal’s cranial anatomy [9,16].

### Identifying Long-Range Projections

With extracellular recordings, cell types cannot be directly identified, but they can often be inferred using features extracted from the shape of their action potential waveforms [17–19]. Most previous work, especially in primates, has relied on the observation that the trough-to-peak width of these waveforms forms a bimodal distribution, with modes thought to correspond to putative excitatory and inhibitory neurons [17]. However, some work in other species, including rodents [20,21], cats [22], songbirds [23], and wallabies [24], suggests that a more detailed analysis of action potential waveforms can potentially isolate contributions from long-range projections.

To do so, we exploit several biophysical phenomena which alter the shape of an action potential as it flows through a neuron, as illustrated in Figure 1A. The opposing dipoles along the axon produce a prominent early positivity; this sharp peak is not present in somatic recordings [25,26]. Additionally, the presence of many rapidly-acting Kv1 channels along the axon causes axonal action potentials to be extremely narrow, even moreso than those produced at the soma of fast-spiking interneurons [27]. Axonal action potentials often include a small and rapidly-repolarizing late peak as well, though its prominence depends on the relative position of the axon and recording electrode [28]. These signals are small, but their magnitude increases with the diameter and branching of the axon [26]. Thus, any axonal signals we detect are likely to arise from larger long-range projections. Consistent with this hypothesis, putative axons tend to have tuning properties consistent with a region’s known afferents [20], such as LGN-like receptive fields in primary visual cortex [22,24]. We pharmacologically validate this conjecture below.

**Figure 1.**
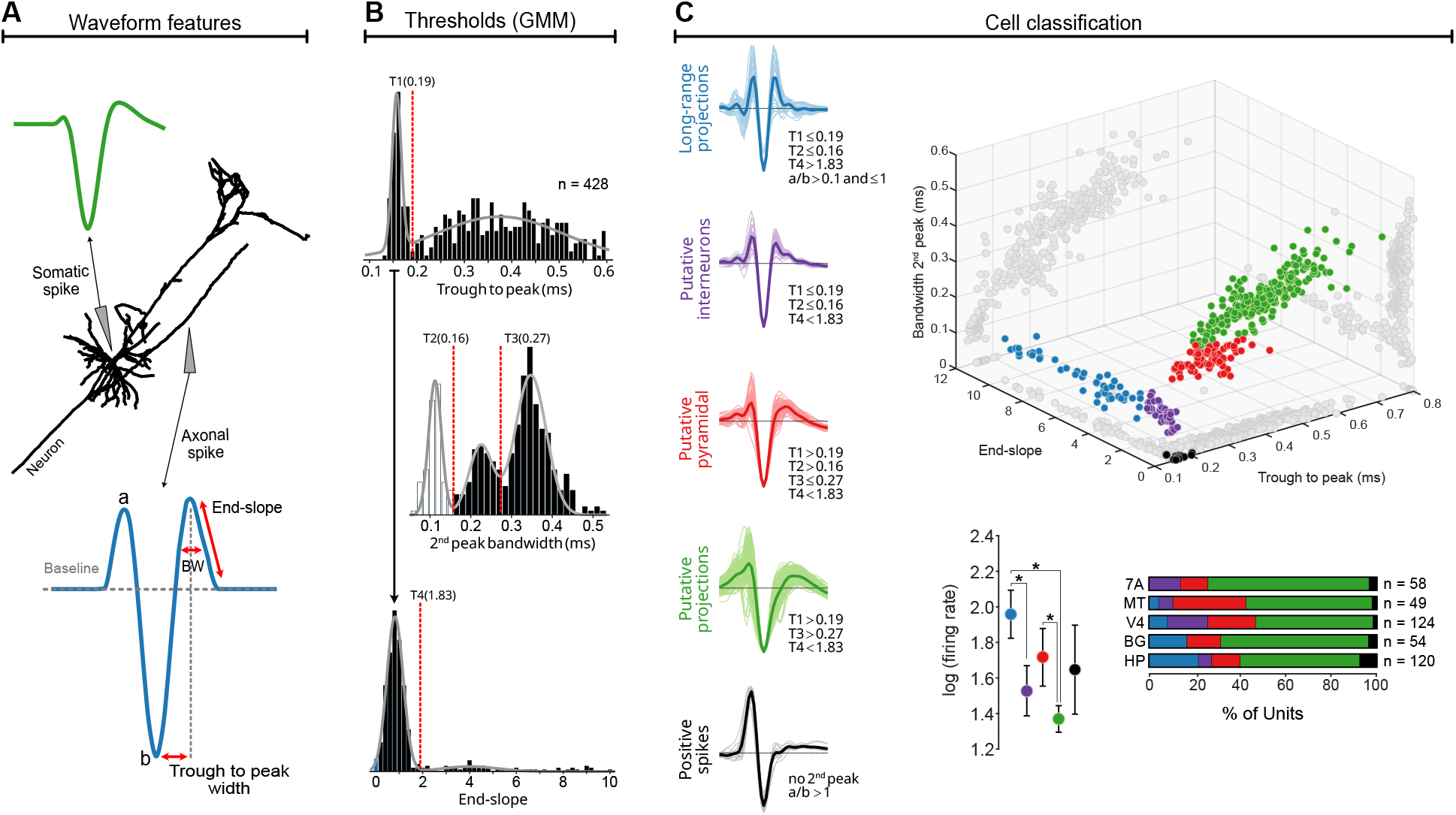
Identification of cell types. A) Sketches of action potentials recorded near the soma (green) and along an axon (blue). Based on the underlying biophysics, signals recorded at these locations have very different shapes. We extracted features capturing these differences, as indicated on the blue waveform. B) Distributions of these features and threshold values (dashed red lines) are shown in the histograms; see *Methods*. C) Based on these features, we divided our data into five cell classes: putative axonal recordings from long-range projections (blue, black), interneurons (purple), and pyramidal cells (red, green). Individual (thin lines) and average waveforms (thick) are shown for each class, with the threshold values shown beneath. As the 3D scatterplot shows, these classes form distinct clusters in the joint feature space. The clusters differ in average firing rate and distribution across brain areas. BW: Bandwidth, BG: Basal Ganglia, HC: Hippocampus. Cell in Panel A adapted from the data of Hallerman et al. [26,30].

These properties should give long-range projections a characteristic shape that is readily distinguishable from other units in primates. We therefore extracted a set of waveform features that capture these biophysical differences, as illustrated in Figure 1A. The a/b ratio and end-slope describe its shape, while the trough-to-peak width and second peak bandwidth (BW) capture the duration of the action potential. Based on the abovementioned biophysical considerations, signals from long-range projections are expected to have large (i.e., rapid) end-slopes and narrow (i.e., short) trough-to-peak width and second peak bandwidths [20,24]. We further filtered candidate waveforms using the ratio of the peak to trough amplitude (a/b in Figure 1B), to verify that each putative long-range projection had the expected shape [20,21,29]. Since the precise value of these parameters may vary across species, electrodes, and recording apparatus, we used Gaussian Mixture Modeling (GMMs) to identify threshold values in a principled and parsimonious way, rather than relying on previously published values.

Using these features (see *Methods*), we identified 49 putative axonal recordings (triphasic long-range projections; blue traces in Figure 1B), 40 putative interneurons (purple), 71 putative pyramidal neurons (red) with broad spikes and 229 more pyramidal neurons with broader spikes (green), as well as 16 putative positive-only axonal recordings (black). When plotted jointly (Figure 1C, right), these clusters occupy very distinct portions of the 3D parameter space and differ in average firing rate and areal distribution. The putative axonal recordings (blue and black), which are of primary interest here, are particularly well-separated from the remaining unit types.

A data-driven approach using non-linear dimensionality reduction [18] led to similar conclusions (Figure S1). Our clusters were in separate parts of the reduced UMAP space and transects through the embedded revealed that arrangement was based on similar waveform features. (Figure S1B). However, due to technical concerns about the interpretation of UMAP spaces [31], we use the clusters generated by our biophysically-motivated features for the remaining analyses instead.

### Pharmacological Validation

Our classification relies on correlations with known biophysical properties, which provides circumstantial evidence that it identifies long-range projections. We performed a separate set of experiments to directly test the origin of these signals with microinjections of muscimol, a potent GABA-A agonist. Muscimol is typically used to produce temporary lesions by silencing neural activity near the injection site [32,33]. It does so by affecting chloride channels that are predominately found on the soma [34]. Although some chloride channels may be found on axons, treating axons *in vitro* with muscimol does not block spiking [35]. Thus, neural activity generated near the injection site in our animals should be silenced by the application of muscimol, but projections arising from distant cells should be unaffected by the drug. If units with triphasic (Figure 1C, blue) and/or positive-going (Figure 1C, black) waveforms persist, this would be strong evidence that they do arise from long-range projections.

Figure 2 shows neural activity recorded before and after muscimol microinjections in two animals. Prior to the injection, we were able to isolate a wide range of units, color-coded as in Figure 1. We next injected 2 μL of muscimol (10 mg/mL) at a rate of 0.05 μL/minute and waited 40 minutes, a period sufficient to produce localized behavioral deficits in behaving animals [32,33]. The waveforms of most previously-identified units became undetectable (red Xs in Figure 2A and Figure 2D) and their firing rates dropped to zero (Figures 2B and 2E). It is unlikely that this simply reflected movement of the probe and/or brain, as few new single units appeared and the multiunit firing rate was also significantly reduced in both animals (11.6 Hz to 1.6 Hz in Figure 2C and 14.9 Hz to 1.9 Hz in, Figure 2F; p<0.03; Wilcoxon Sign-Rank test).

**Figure 2.**
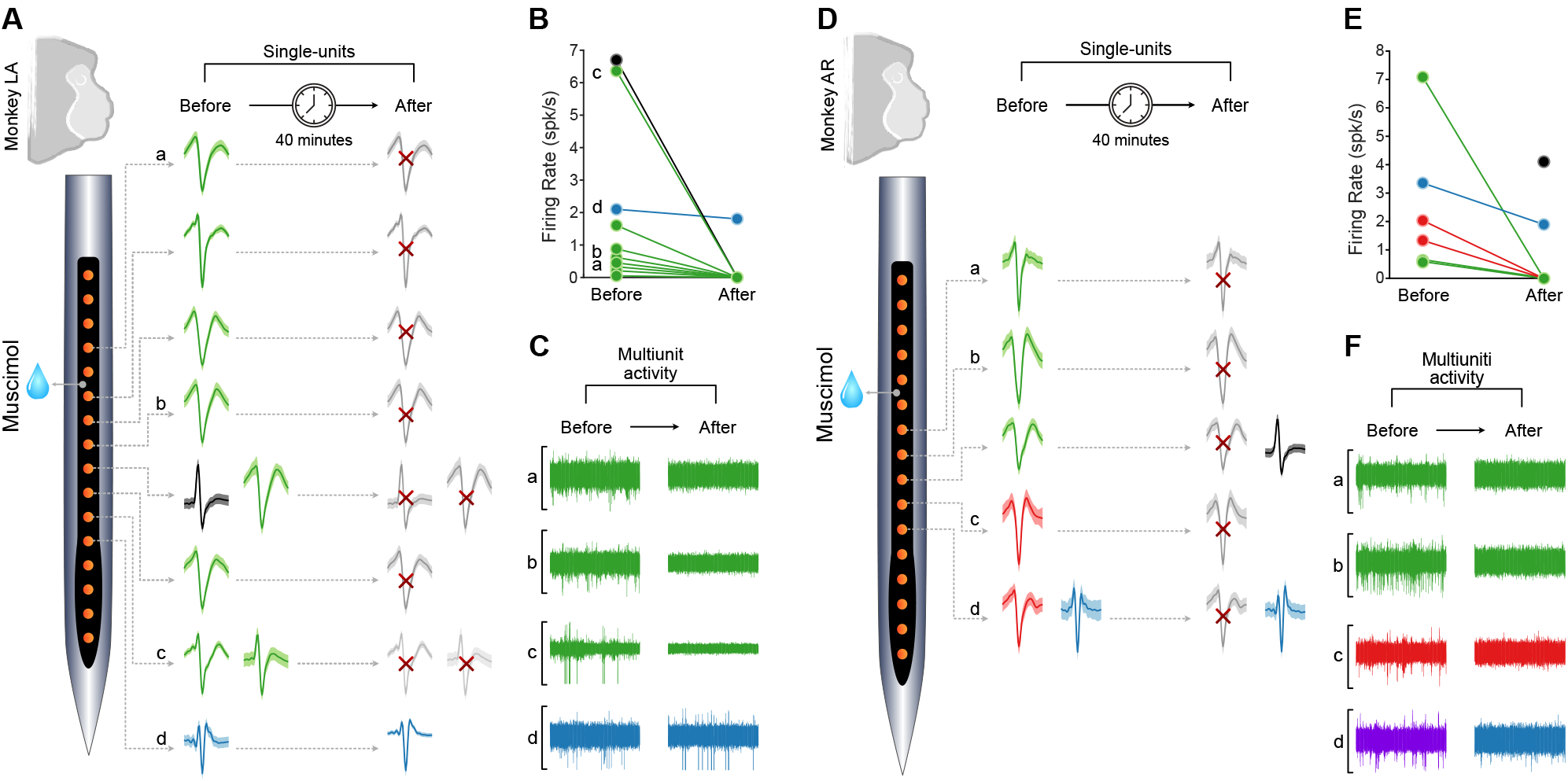
Only signals from putative axons persist after local activity was pharmacologically silenced. A) Averaged extracellular waveforms (± one standard deviation) recorded before (left) and 40 minutes after (right) injecting muscimol, a potent GABA agonist. Waveforms are color-coded as in Figure 1 and are shown based on their relative locations on the microelectrode array. Red Xs indicate neurons that were no longer present after injection. B) Average firing rates for isolated single-units before and after muscimol injection. C) High-pass filtered signal on four example channels, indicated in Panel A, showing the lack of multiunit activity as well. D-F), Same as A-C, but in a second animal.

Nevertheless, we were still able to isolate a small number of units. This includes two putative long-range projections (Figure 2A and 2D, blue), which were present before muscimol injection and maintained their spiking activity afterward. In one case, this was accompanied by the silencing of a broad-spiking unit that was previously isolated on the same channel (Fig 2D, red). Finally, we also isolated a third unit (Monkey AR) which was not present before the injection, but that had a waveform consistent with an long-range projection. Due to the geometry of our electrode arrays, our recordings come from within 1.2 mm of the injection site. Since the drug is expected to diffuse ∼2 mm during this time [36], these data demonstrate that these units arise from more distant sources, or in other words, constitute long-range projections.

### tACS Preferentially Entrains Projections

Having established that we can identify long-range projections via their extracellular action potential waveforms, we next asked how they are affected by tACS. A large body of work now suggests that the principal effect of tACS is to change spike timing, rather than rate [12,14,37–39], especially rhythmic activity occurring at the same frequency as the stimulation. We measured these effects by calculating phase-locking values (PLVs; see *Methods*), a measure of entrainment between spikes and an oscillation. A PLV value of 1 indicates that spiking occurs at exactly the same point in each cycle of the oscillation; 0 indicates that the neuron fires at random without any regard to the oscillation. We separately calculated PLVs during stimulation and baseline/sham conditions to derive a ΔPLV value that indicates the effects of stimulation. Note that ΔPLV values can be less than zero if a neuron fires rhythmically during baseline and stimulation disrupts this rhythm; this is a common outcome during tACS [14,40].

Figure 3A shows the ΔPLV for all 405 classifiable units, colored by cluster. Putative long-range projections had a larger ΔPLV (median: 0.025, 95% CI: [0.01, 0.06]) than putative interneurons (median: 0.007, 95% CI: [-0.01, 0.03]), putative pyramidal neurons with broad (median: 0, 95% CI: [-0.010, 0.013]), or very broad spikes (median: 0.0019, 95% CI: [0, 0.013]). These differences were all statistically significant (all *p* < 0.03; Wilcoxon Rank Sum tests). There was, however, no significant difference between triphasic and positive-only units (median: 0.009; 95% CI: [0, 0.054], p>0.25, Wilcoxon Rank Sum tests), consistent with the idea that the two classes both represent long-range projections. Since these data were collected using multiple electric field strengths, we repeated this analysis after normalizing the data in terms of ΔPLV per V/m, but this led to identical conclusions (Figure S2).

**Figure 3.**
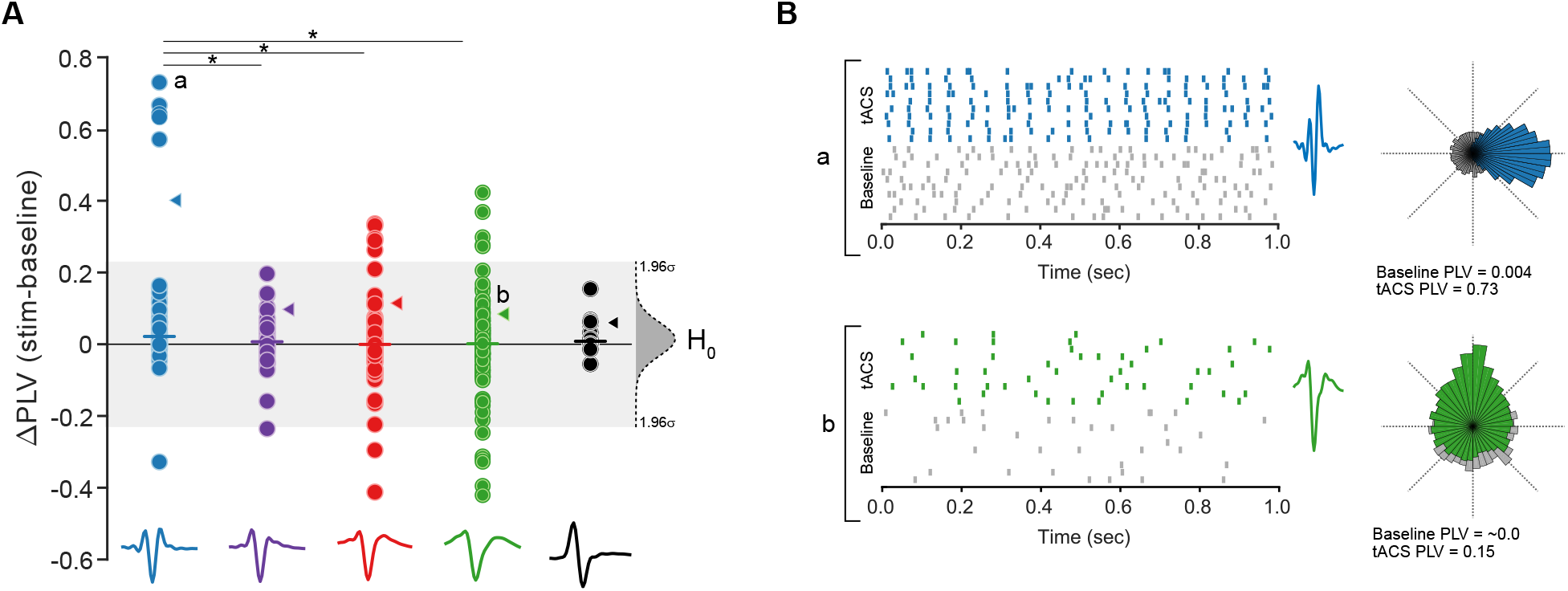
Putative long-range projections are more strongly affected by tACS. A) Changes in phase-locking values for each of the 405 classifiable neurons in our dataset, colored by cell class (as in Figure 1 and 2). Horizonal lines indicate the medians for each class, arrowheads indicate the 90^th^ percentile. The grey region indicates the 95% confidence interval around the overall mean. B) Raster plots showing the activity of two example neurons: a putative axonal recording from a long-range projection (labelled a in Panel A) and a putative pyramidal cell (labelled b). Spiking activity during sham is shown in grey, activity during tACS is shown in color. Polar histograms showing the phase of spiking relative to the stimulation (color) and local field potential (grey) demonstrate how phase-locking changes; PLV values are shown below each histogram. * indicates p < 0.05

These data demonstrate that a typical axonal unit is more strongly entrained by tACS than other cell types. However, as Figure 3A shows, a handful of units seem to respond unexpectedly strongly to tACS. Twenty-seven units had ΔPLVs outside the 95% confidence interval of the entire dataset, including five points that were each more than 5σ away from the mean. Notably, these cells were all classified as long-range projections. In these cells, tACS produced activity that was extremely well-synchronized to the tACS waveform, as shown in the example cell of Figure 3B (top panel, blue). This contrasts with the relatively modest effects which typically occur during stimulation; the neuron in the bottom panel of Figure 3B has a ΔPLV of 0.15, a typical effect of tACS at 1 V/m [12,38]. These points are even more striking because 3 of the 5 units were recorded during our weakest stimulation condition (0.3 V/m).

These extremely responsive units appeared to occur more frequently within certain clusters. To test this hypothesis more formally, we used quantile regression [41], a technique used to analyze differences in the tails of distributions rather than their central tendencies. Here, we asked whether the 90^th^ percentile ΔPLV differed between groups. Specifically, we determined if the data was more parsimoniously described as a single distribution, combing all 406 units together (i.e., a “null model”) or as five separate distributions reflecting the predicted class of each unit. The latter model had a dramatically better fit (ΔAIC = -29.6), even when accounting for the additional parameters. Moreover, most of this improvement could be captured by a simpler model which considered only whether a unit is a putative long-range projection (blue, ΔAIC = -25.4). In the full model, coefficients for all classes other than putative long-range projections were negative, and, with the exception of the putative (very broad) pyramidal cells, significantly so (p < 0.05). Since these extreme data points do not affect the median, which is robust against outliers, this analysis highlights a separate aspect of the data. Thus, on average, long-range projections are more entrained by tACS and are overrepresented among the cells that are extremely affected by it.

## Discussion

Here, we show that units corresponding to long-range projections can be reliably identified via the shape of their extracellular waveform in non-human primates (Figure 1). This identification is consistent with their known biophysical properties and was confirmed pharmacologically via microinjection of muscimol (Figure 2). We use these data to demonstrate that long-range projections are especially susceptible to electrical stimulation (Figure 3), which may have important implications for the design of therapeutic and scientific applications of brain stimulation.

## Challenges for focality

Most brain stimulation experiments are motivated by the idea of changing one localized activity pattern and allowing the results of that perturbation to influence the rest of the brain and ultimately behavior (Figure 4A). Considerable effort has thus been devoted to developing increasingly more focal stimulation techniques, using personalized montages [9], multiple electrodes [7], and interference patterns [5,8,42] to limit the spread of the electromagnetic field during non-invasive brain stimulation. These techniques do better control the flow of current into the target. However, if that stimulation preferentially affects long-range projections, its main effect will be to modulate inputs and outputs from that target, rather than local processing (Figure 4B).

**Figure 4.**
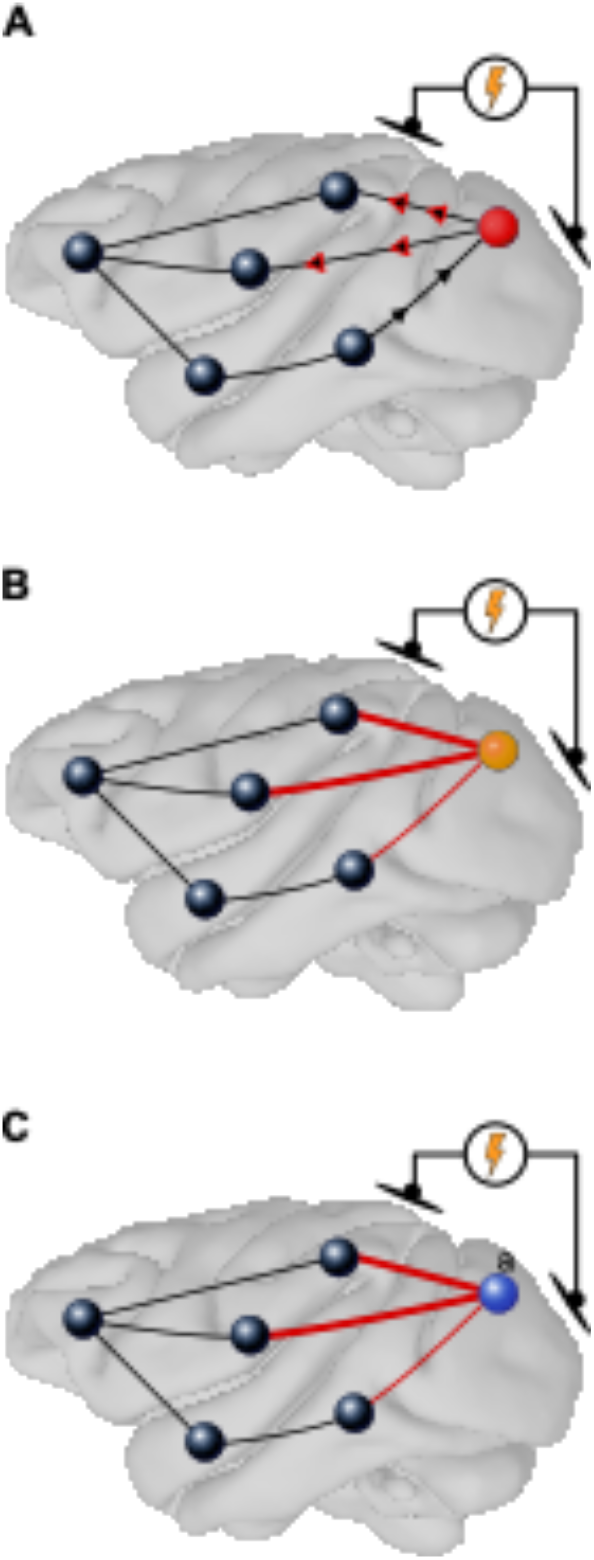
Schematic depiction of stimulation effects. A) Stimulation is traditionally expected to affect local computations in the targeted region and the results of that perturbation are passed onto other brain regions (arrowheads) via its outputs (lines). B) Our data suggests that stimulation not only modulates local processing but strongly affects inputs and outputs to the target area. C) In an extreme case, the local and long-range effects may even be qualitatively different, with local activity desynchronized but long-range connections entrained to the stimulation.

Because stimulation competes with other endogenous influences for control over spike timing [14,40], this may even produce categorical differences between the local and long-range effects. Weak stimulation of local neurons may desynchronize them without imposing a new rhythm, while still entraining long-range projections (Figure 4C). This same competition explains why the median PLVs are near zero for some cell types (Figure 3). We stress this does not imply that tES is ineffective overall: forty percent of the units in our data set had individually significant changes in PLV. Instead, the sign and direction of each cell’s outcome determined by the relationship between the stimulation and the neuron’s ongoing activity [14,43]. Predicting the net effects of stimulation may therefore require considering how both local and long-range projections couple to each other as well as the stimulation [40].

### Why does white matter stimulation work?

Several recent studies have reported that long-range projections, in the form of white matter tracts, may be a promising target for invasive [44–46] and non-invasive brain stimulation [47– 49]. For example, during a verbal working memory task, stimulation delivered near white matter was 28% more effective than stimulation delivered farther away [44]. Similarly, it has been suggested that white matter might be a more effective target in treating a wide range of conditions, including depression [45] and obsessive-compulsive disorder [50]. This relates to the growing appreciation that many psychiatric and neurological disorders can be characterized as “disconnection syndromes”, in which connectivity between regions is disrupted, rather than as impairments in local brain function[51].

The efficacy of these stimulation protocols is generally attributed to white matter’s functional roles: the signals it carries may be more tied into distributed brain networks that mediate behavior [44,52,53]. Alternately, it may be easier to design appropriate interventions for the lower dimensionality of the signals they carry [52]. Our results do not directly argue against these interpretations, but they do suggest that they need to be carefully tested to avoid a trivial confound: the same stimulation will produce larger effects on long-range projections.

Other studies have argued for the importance of multi-site or “network” stimulation that is specifically tailored to affect connectivity between areas. These protocols often seem to work [54,55] and our results suggest that, happily, they may be relatively easy to achieve. It may be possible to stimulate long-range projections between areas with a single pair of electrodes if the location and orientation of the induced electric fields is chosen carefully. In this view, methods for generating more focal electromagnetic fields may be especially valuable because they permit one to precisely target a particular set of projections.

### Reliability of axonal identification

These conclusions depend on an accurate identification of axonal vs. somatic activity. Our results are limited insofar as they rely on circumstantial evidence, the shape of the extracellular action potential, rather than direct labelling or measurements. Unlike prior attempts to identify excitatory and inhibitory neurons via spike width, the muscimol experiments (Figure 2) provide rigorous external validation of our clustering. In other animal models, functional properties of units with similar waveforms provide additional evidence that they arise from long-range projections. For example, triphasic units in Area 17 of the cat and wallaby lack orientation tuning, as would be expected of thalamic inputs [22,24]. Our simple task did not permit us to analyze the functional properties of these units in macaques, but this will be a rich direction for future work.

Much of the prior work investigating axonal activity in primates relies on technically challenging approaches like opto-tagging [56] or antidromic activation [57] to identify projections, and can provide detailed information about the source and target of each neuron. Likewise, the use of ultra-high density electrodes has recently been proposed as a method for identifying axonal projections by collecting additional information about the electrical “footprint” of each unit [58], which is thought to be smaller for axons [26]. It may be possible to approximate this feature by slightly moving a standard electrode array. Even without doing so, our results suggest that putative axons can be readily identified from standard recordings, even within existing data sets. This only requires modifying spike-sorting protocols so that atypical waveforms are not excluded early in the process.

### Alternate Interpretations

One might be concerned that axonal projections are not intrinsically easier to entrain but simply arise from areas which receive more of the stimulating current. Indeed, this argument has been advanced to explain all stimulation effects occurring below the skin or cortical surface [59] (but see [60]). If this trivial explanation were true, we would expect to see the strongest effects in units that were recorded both near the focus of stimulation and at the cortical surface. This was not the case. None of our putative long-range projections were recorded from Area 7A (Figure 1C), where tACS produced the strongest electric fields in our dataset [15]. Instead, three of the 5 extremely entrained units (Figure 3, blue) were recorded in deep structures; the other two were recorded in Area MT. Even if the cell bodies of these projections were located on the cortical surface, this alternate hypothesis predicts that they should be as affected as the local circuitry recorded elsewhere.

During suprathreshold stimulation, it is well-known that stimulation of the axon—and specifically the axon initial segment (AIS)—produces the largest effect for a given bolus of current [61]. However, our stimulation is much more diffuse than the size of an AIS and is not specifically targeted at it. Thus, enhanced stimulation of the AIS alone is unlikely to explain our results, especially because every neuron, not just long-range projections, has an AIS. However, other properties of axons may make them particularly susceptible to electric fields: in vitro experiments have found that electric fields preferentially polarize axons[62,63] and alter the probability of synaptic release[62]. These effects may increase with the size of the neuron, producing larger effects on larger long-range projections. Alternately, it is usually assumed that electric fields are approximately uniform on the scale of a single neuron. However, long-range projections in primates can be many millimeters long and thus may be affected by electric field gradients as well as morphological features.

Owing to their larger size, long-range projection neurons appear to be more affected by tACS on average. We speculate that extremely strong effects occur in a subset of these neurons which are both large and optimally aligned with the electric field [64,65]. We did observe some relatively large effects in the very broad spiking class (green, Figure 3A) and it is possible that some of these units may be somatic recordings from outgoing projections: neurons projecting from V1 to V2 have broader spikes, but this criteria alone was not enough to conclusively identify them [66].

Taken together, our results suggest that brain stimulation protocols must consider a combination of local effects on the targeted structure as well as potential modulations of its inputs and outputs. This may complicate interpretations of behavioral effects but suggests that it may be possible to create broad changes across networks of areas with relatively simple methods.

## Acknowledgments

We thank Dr. Fernando Chaurand, Julie Coursol, Cathy Hunt, and Jessica Hutta for outstanding technical assistance. This work was supported by grants from NSERC (DH-2022-00476) and the CIHR (202104PJT-461642).

## Methods

This paper reanalyzes data from several of our prior publications. Critical details of the methods are described below. Interested readers are referred to Krause and colleagues [14] for additional details and validation of the brain stimulation techniques and Laamerad [33] for details of the inactivation methods.

The brain stimulation data were originally reported in several prior studies [12–15]. We limited our analysis to epochs where 5, 10, or 20 Hz conventional low-frequency tACS was applied. In particular, we excluded the conditions using other stimulation modalities in Vieira and colleagues [15]. The combined dataset consists of recordings from 428 single units from four rhesus macaques (two male, two female) receiving tACS. The pharmacological experiments were previously reported in in studies focused on the behavioral consequences of inactivation [32,33], rather than the residual activity described here. Those experiments collected data from two female monkeys.

### Ethics Statement

These experiments were overseen by the Montreal Neurological Institute’s Animal Care Committee (Animal Use Protocols #1870 and #5031), using additional guidelines from the Canadian Council on Animal Care and the *Weatherall Report on the Use of Non-Human Primates in Research*. Day-to-day operations were supervised by the Institute’s veterinarians and trained animal health staff. Animals received daily and varied environmental enrichment, both in their home cages and through regular access to a large play arena, where they had visual and tactile access to conspecifics. Positive reinforcement techniques were used to train animals to transfer from their home/play cages to their primate chairs and to perform behavioral tasks.

### Animal Preparation

We performed a series of scans and surgeries to prepare animals for these experiments. For each animal, we initially acquired a set of T1 and T2w MRIs (0.6mm isotropic voxels). These scans were used for electric field modelling and presurgical planning. Next, we attached a titanium head holder used to stabilize the animal during electrophysiological recordings. Animals were then rescanned with a fiducial marker attached to the headpost to provide reference points for subsequent neuronavigation. Once animals were then familiarized with the laboratory, head restraint, and the behavioral task, we implanted a recording cylinder (Crist Instruments, Hagerstown Maryland) to provide access to target areas. This included the basal ganglia and hippocampus (one animal each) and multiple visual areas in two other animals. This was done under guidance from a frameless stereotaxy system (Brainsight, Rogue Research; Montreal, QC) with submillimeter resolution. Electrode positions were verified via MRI or CT in three of the four animals.

During MRI scans and surgical procedures, animals were initially sedated with an intramuscular injection of ketamine (∼10 mg/kg), followed by inhalational isoflurane (∼1-2%). These were adjusted to achieve a stable anesthetic plane, as per McGill’s Standard Operating Procedure #115. A qualified veterinarian oversaw post-procedure treatments and prescribed any necessary analgesia and antibiotics. However, all data reported here was collected from awake, behaving monkeys who had fully recovered from the procedures.

### Neurophysiological Recording

We made daily-acute recordings through the recording chambers implanted in each animal. We initially penetrated the dura with a 22-gauge stainless steel guide tube, then inserted 32 channel linear arrays (V-Probe; Plexon, Dallas, Texas, USA). Each site on the array had an impedance of 200—400 kΩ; adjacent sites were separated by 150 μm. During muscimol experiments, 16-channel V-Probes with an integrated fluid channel were used instead. Arrays were then lowered into the target structures by a computer-controlled microdrive (NAN Instruments, Nazareth Illit, Israel), using depths derived from the imaging and neuronavigation system. These were generally consistent patterns of activity we observed while lowering the electrode.

During stimulation experiments, signals were recorded from these electrodes with a Ripple Neural Interface processor (Ripple Neuro; Salt Lake City, Utah) and sampled at 30,000 Hz, with 16 bits per sample and 0.50 μV per least-significant bit. The amplifier’s linear range spans ±12 mV and we adjusted grounding, amplifier, and stimulation settings to ensure that the recording system was not saturated and remained in that range. The raw signal was initially bandpass filtered between 0.3 – 7,500 Hz by hardware filters in the recording hardware. During muscimol experiments, data was instead collected with an Intan Technologies RHD system but was processed similarly.

Signals were further processed offline to extract spiking activity. The recorded signals were always bandpass filtered between 700-5000 Hz and spike thresholds were set at ±3σ, robustly estimated via the median absolute deviation. Snippets around each threshold crossing were extracted and clustered with UltraMegaSort2000, a *k*-means overclustering spike sorter[67], followed by manual review to ensure they had a consistent shape, clear refractory period, and good separation in PCA space. We have extensively tested this pipeline against potential stimulation artifacts [12,15].

### Brain Stimulation

Our stimulation procedures were designed to closely mimic human neuromodulation and therefore used commercially available equipment marketed for human use. Stimulation was applied to the scalp using 1 cm (radius) silver/silver chloride electrode (PISTIM; Neuroelectrics, Barcelona, Spain). Each electrode was coated with a conductive paste (SignaGel; Parker Labs Fairfield, New Jersey) and attached to the intact scalp with a biocompatible silicon elastomere (KwikSil; World Precision Instruments, Sarasota, FL]). Electrode impedances were typically 1-2 kΩ and never exceeded 15 kΩ.

To place the electrodes, we used finite-element modelling to identify suitable scalp locations for stimulating our target structures, using either SimNIBS [9] or a custom pipeline [16]. Stimulation currents were limited to ±1 or 2 mA and produced electric fields of 0.3 to 1 V/m. Specifically, electric fields of 0.3 V/m were produced in the hippocampus and basal ganglia, 0.9 V/m in V4d, 0.6 V/m in 7A, 0.5 V/m in MT, and 0.4 V/m in V4v. These values span the range expected to occur in human use [68,69].

For two animals, electrical stimulation was delivered through two NeuroConn DC-STIMULATOR PLUS devices (NeuroConn, Ilmenau, Germany). The device was operated in “remote mode”, where each converted a voltage input into a constant current output (1V = 2 mA peak-to-peak, or ±1 mA). The voltage input was provided by a B&K Precision Model 4053B waveform generator, which was programmed to emit sine waves. In the other two animals, stimulation was delivered by a StarStim8 system (Neuroelectrics, Cambridge MA), which was controlled via its MATLAB interface.

### Muscimol Injection

Microinjections of muscimol were used to validate our classification of units as long-range projections. Muscimol (Sigma-Aldrich, St. Louis, MO) was diluted to a final concentration of 10 mg/mL in sterile saline and passed through a 0.22 μm filter. We recorded neural activity at the beginning of each session to verify that the probe was inserted and functioning correctly. Once units were isolated, we injected 2 μL of muscimol (10 mg/L) at a rate of 0.05 μL/minute using a 5 μL glass syringe (Hamilton, Reno, NV) driven by an infusion pump (Chemyx Fusion 100; Chemyx, Dallas, TX) and connected to an integrated fluid channel inside the multielectrode array. were used instead. Fluid exited between the 5^th^ and 6^th^ contact, as shown in Figure 2. After waiting 40 minutes for the drug to diffuse, we again recorded neural activity. During this time, muscimol is expected to diffuse approximately 2mm [36], which is farther than the span of our array (up to 1.2 mm from the exit point).

Injection of muscimol caused a dramatic decrease in neural activity, as discussed above, but no changes were observed after a saline (vehicle) injection. We interpret this as demonstrating that these decreases are due to activation of chloride channels, rather than fluid pressure or other confounds. In one experiment, a putative long-range projection did disappear. These data are difficult to interpret. It could reflect a projection from a neuron whose soma was also in the inactivated region. However, because micromovements of the brain and/or probe over the hour-long experiment could also lead to loss of isolation; axons, being smaller than the soma, have a smaller electrical “footprint” and are thus harder to stably record[20]. However, it is unlikely that this effect could account for the loss of all other units and the reduction in multiunit firing rate.

### Quantification and Statistical Analysis

In non-human primates, practical and ethical considerations limit the number of animals from which we can collect data. This study reanalyzes existing data from four animals and the critical comparisons in this paper are either within-cell (ΔPLV) or across cell types, rather than within animals. Thus, the animal is not the relevant unit of analysis in this work [70], which also makes it impossible to distinguish sex differences from other individual differences.

We did not carry out a formal power analysis since little is known about this phenomenon, especially in primates and the present work reanalyzes all of our available data. Based on our prior work with tACS, changes in phase locking can be reliably characterized with 1.5 to 5 minutes of data. Other work has found that the strength of entrainment, though not necessarily the preferred phase itself, is rapidly established after stimulation onset and remains constant on these timescales[71]. To the extent that data limitations hinder our ability to detect weak entrainment, we would expect this to bias our analysis away from finding differences in entrainment between cell classes.

Where possible, statistical analyses used nonparametric tests to avoid distributional assumptions, and all reported values are two-tailed. Confidence intervals for the medians were calculated using the formulae of Campbell and Gardner[72]. Data were analyzed using MATLAB (The MathWorks), R (Foundation for Statistical Computing, Vienna, Austria), and the quantreg package [41].

### Analysis of Extracellular Waveforms

Extracellular action potential waveforms were initially extracted by the spikesorter, as described above. Each unit’s waveforms were then up-sampled 400x with a cubic spline, aligned to their troughs, and averaged. The features described above were then extracted from the average extracellular waveform.

For each feature, we identified clusters and cluster boundaries by fitting Gaussian mixture models (GMMs) to each feature histogram. We considered possible mixtures containing from 1-10 components and chose the most parsimonious on the basis of Bayesian Information Criteria. To avoid local minima, the modelling procedure was repeated 100 times with randomly-chosen starting points. Cut-points were derived from inflection points in the best-fitting model. This procedure is very similar to the one used by Traintio and colleagues [19].

The trough-to-peak width was initially used to divide the population into narrow- and broad-spiking units. Consistent with prior work [15], our analysis suggested the existence of two clusters, separated at a threshold of 0.19 ms (Figure 1B, dotted red lines on histograms). However, the narrow group likely includes both somatic recordings from putative interneurons, which also have rapid potassium channels, as well as recordings from the axons of long-range projections [73]. Thus, we further divided these cells based on the end-slope parameter. Our GMM analysis also suggested the existence of two clusters in the end-slope data, with the steeper end-slopes likely corresponding to long-range projections. We additionally required these units to have an initial peak-to-trough ratio of at least 0.1 to ensure they had a triphasic shape [20,21].

The broad-spiking units were also subdivided into clusters, this time via the second peak bandwidth. Here, the GMM found three distinct clusters. The narrowest bandwidth cluster (white bars) was not informative, as it heavily overlapped with the narrow-spiking units identified above. The other division at 0.27 ms was used to split units into broad and very broad clusters, a division seen in other work [19] and which may correspond to somatic recordings from local (broad) vs. outgoing projection (very broad) neurons [66]. Finally, we identified a fifth cluster of positive-going waveforms that lack a clear second peak and thus usually have a peak-to-trough (a/b) ratio larger than one. The end-slope could not be computed for these units, so we consider them as a separate cluster, but one that likely also reflects axonal activity [28]. The vast majority of units (95%; 405/428) units could be assigned to a single cluster based on this procedure. We excluded the remaining 21 neurons from the rest of our analysis.

As an alternate approach, we performed an unsupervised analysis of our dataset using WaveMap [18], a recently developed pipeline that combines non-linear dimensionality reduction technique (UMAP) with graph clustering (Louvain community detection). In brief, raw waveforms were non-linearly embedded in a high dimensional space. Louvain clustering was performed in the high dimensional space, after which the data were projected into two dimensions for visualization. The results of this unsupervised method closely resemble our other clusters (Figure S2B).

### Spike Timing Analysis

We quantified neural entrainment by calculating for each cell the pairwise phase consistency (PPC) value, a measure of the synchronization between the phase of a continuous signal (like the LFP or tACS waveform) and a point process like single-unit spiking activity[74]. Because spikes can introduce small but detectable changes in signals recorded from the same electrode[75], we compared spikes on one channel with the continuous signal from an adjacent channel 150 μm away. By using a local signal, rather a copy of the stimulator output, referencing conditions remain constant across conditions and any physiological distortion of the signal is incorporated into our measurements[76]. During baseline, the continuous signal was the LFP, filtered in a ±1 Hz band around the stimulation frequency used in the rest of the experiment. The Hilbert transform was then used to derive the instantaneous phase of the continuous signal at the time of each spike. These were used to calculate PPC values, which have several desirable statistical properties: notably, it is unbiased and unaffected by the number of spikes. Since PPC values are rarely reported in the literature, we transformed them to phase locking values (PLV) for display by taking their (positive) square root. The resulting PLVs are also equivalent (under some simplifying assumptions) to spike-field coherence, another oft-used metric of spike-LFP coupling.

## Supplemental Figures

**Figure S1.**
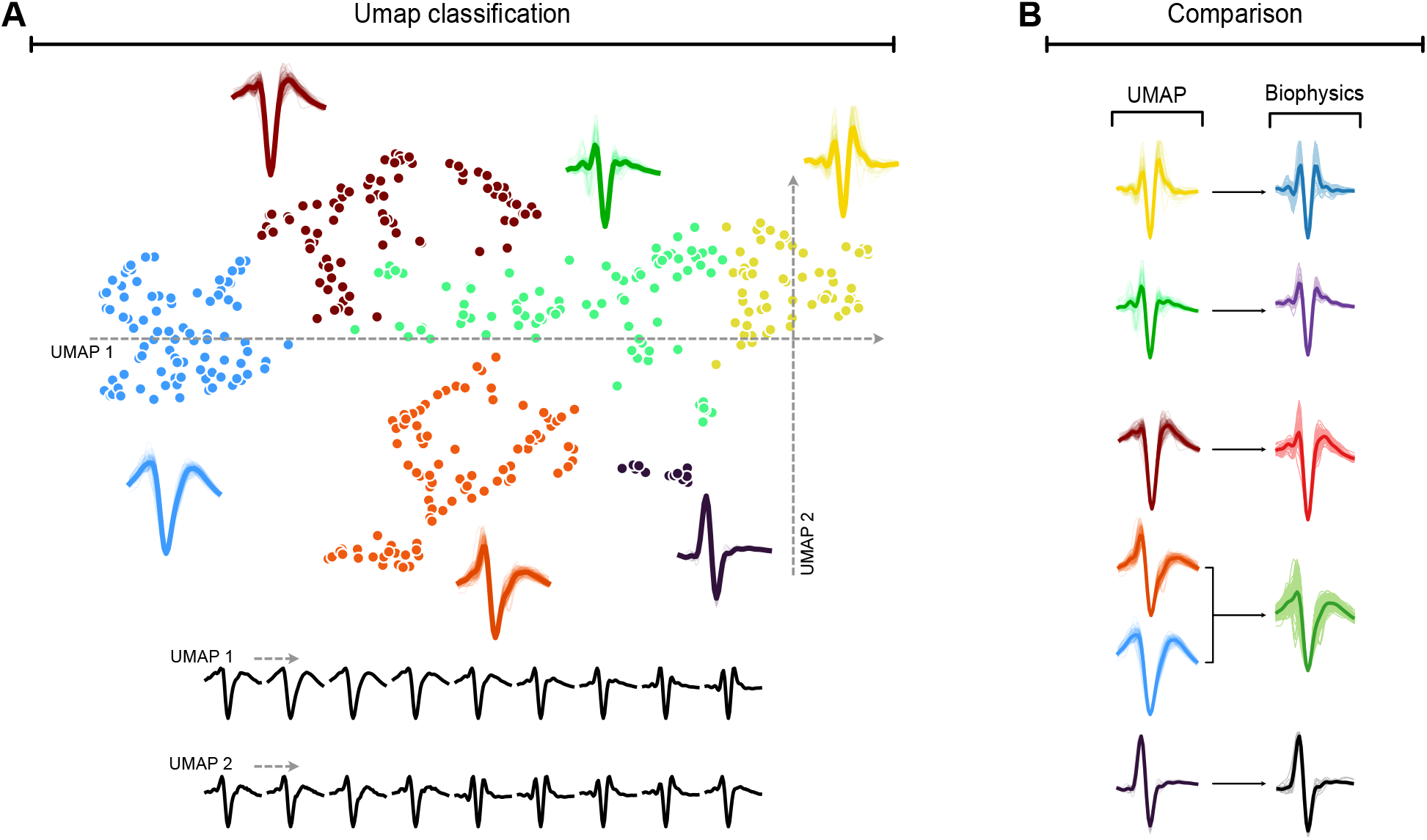
Data-driven classification of cell types. A) Amplitude-normalized waveforms were non-linearly embedded in a high dimensional space via UMAP, clustered, and projected into two dimensions for visualization. Colors indicate the results of the clustering and insets denote the average extracellular waveform for each cluster. Transects through the embedding space revealed that the relevant dimensions resemble those captured by our features: the horizonal dimension appears to encode trough-to-peak width and ratio, while repolarization slope varies along the vertical dimension. B) These clusters map onto the biophysically-informed classification used in the main text.

**Figure S2.**
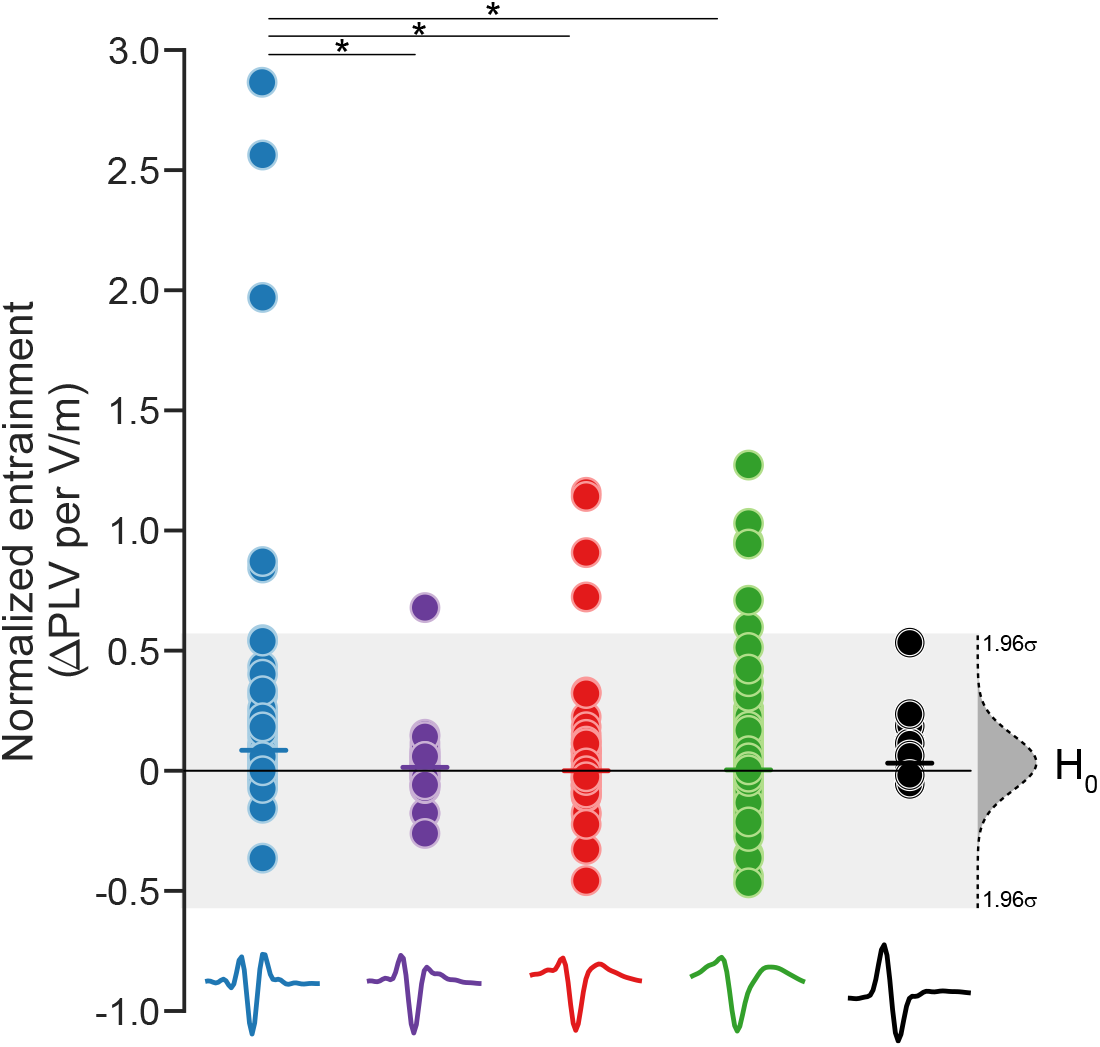
Neural entrainment adjusted for electrical field strength. Since data were collected at different field strengths (see *Methods*), we normalized the ΔPLV values in Figure 3 by the predicted field strength delivered in each experiment, under the assumption that changes in PLV are proportional to changes in field strength [38]. This demonstrates that outlying values are not driven by extremely strong stimulation.

